# Relation between executive functions and polymorphisms in COMT, MAO-A, HTTLPR, SLC1A1 and HT2A in a sample of children with Obsessive Compulsive Disorder

**DOI:** 10.1101/452193

**Authors:** Jorge Enrique Avila Campos, María Cristina Pinto Dussan, Ángela María Polanco Barreto, Esneyder Manuel Guerrero, Rafael Antonio Vásquez Rojas, Humberto Arboleda Granados

## Abstract

**Background:** Obsessive compulsive disorder (OCD) has a complex etiology related to multiple neuropsychological factors. OCD is associated with several candidate genes but results are discordant. The objective was to explore the association between five polymorphisms related to neurotransmitters, the risk of an OCD diagnosis and the performance in four executive functions tests done with Colombian patients diagnosed with this condition.

**Methods:** 63 patients and 65 controls matched by gender and age were genetically analyzed. For the study of the relation between cognitive function and phenotypes, a subsample of 33 patients and 31 controls was used. The Stroop test, Wisconsin Card Sorting Test (WCST), Tower of London and Trail Making Test (TMT) for executive function assessment were applied and the SNPs analyzed were: COMT (rs4680), MAO-A (rs6323), HTTLPR (rs25531), HT2A (rs6315) and SLC1A1 (rs301434).

**Results:** Differences in the conceptualization of the WCST test (p = 0.023) and Stroop interference score (p = 0.041) between cases and controls were obtained. After analyzing the relationship between genotypes and sub-scores of the tests, associations between the presence of MAO-A, SLAC1A1, HTTLPR and HT2A alleles and tests sub-scores were found.

**Discussion:** This characterization of children with OCD is a new field of work in Colombia and one of the first works performed in Latin America. The sample size and the number of polymorphisms analyzed in this population should be increased.

---

According to APA (American Psychiatric Association. & American Psychiatric Association. DSM-5, 2000 and 2013), the obsessive compulsive disorder (OCD) is an entity characterized by anxiety and the presence of obsessions and/or recurring compulsions. Miguel et al. (2005) state that variability in the phenotypic expression has led to the hypothesis that OCD is a heterogeneous disorder that obscures clinical findings, the natural history of the disease, the response to treatment and the search for related genes. Current research on OCD is directed to describe such heterogeneity in order to improve its clinical and physiological classification by including molecular and anatomical aspects (Browne et al., 2014; Cappi et al., 2016; Grunblatt et al., 2014; Katerberg et al., 2010; Mataix et al., 2008; Orefici et al., 2016; Taylor, 2016; van de Vondervoort et al., 2016).

Clinically, in addition to disorder associated symptoms, OCD patients have a performance in neuropsychological tests that allow defining cognitive characteristics. For some authors, the main difficulty for OCD patients is their inability to inhibit ideas and behaviors (Chamberlain et al., 2005; Thomas et al., 2014). For other authors, the dysfunction for these patients is related to executive functions in general (Abbruzzese et al., 1995; Bishara et al., 2010; Kuelz et al., 2004; Lewin et al., 2014; Olley, Malhi, & Sachdev, 2007; Rao et al., 2008). In Latin America, children with OCD have not been characterized but a general dysfunction in executive functions is expected.

Regarding related biomarkers, reports show a clear genetic transmission; however, how such transmission occurs and the involved genes are almost unknown and studies are contradictory (Browne, 2014; Miguel et al., 2005 Nestadt, 2010; Stewart, 2013).The diversity of the samples generates variability in the disorder, so the identification of genetically valid subgroups is necessary for research and, thus, discover consistent genes that allow recognizing pathological mechanisms and developing more effective treatments (Nestadt, Samuels, et al., 2000; Nestadt et al., 2010)

In association studies, reviews, meta-analyzes and GWAS, a list of genes correlated to OCD are reported (Nestadt et al., 2010; Nicolini et al., 2009; Pauls, 2010.) Nevertheless, five genes related to neurotransmitter systems were particularly important for this study: the gene for catechol-O-methyltransferase (COMT) (Azzam & Mathews, 2003; Denys et al., 2006, He et al., 1995; Karayiorgou et al.; Karayiorgou et al., 1999; Nicolini et al., 2009; Wang et al., 2009; Zinkstok et al., 2008) the regulator gene for the monoamine oxidase-A enzyme (MAO-A) (Hemmings et al., 2003; Karayiorgou et al., 1999; Liu et al., 2013; Weyler et al., 1990). For serotonin, genes for 2A serotonin receptor (5HT2A) and the serotonin transporter (5HTTLPR) were considered (Dickel et al., 2007; Graff-Guerrero et al., 2005; Lin, 2007; Nicolini et al., 2009; Rocha et al. 2008; Saiz et al., 2008; Wendland et al., 2007). And the gene encoding the glutamate transporter (SLC1A1) (Arnold et al., 2009; Arnold et al., 2006; Bailey et al., 2011; Kanai and Hediger 2004; Samuels et al., 2011; Shugart et al., 2009; Stewart, Mayerfeld, et al., 2013; Veenstra-VanderWeele et al., 2001; Rosenberg et al., 2004).

Progress in diagnosis has allowed finding that the disorder has a worldwide prevalence between 1% and 4% in adults (Angst et al., 2004) and between 1% and 2.3% in children (Kalra & Swedo, 2009).There are no clear data on the prevalence of OCD in Colombia (MinSalud, 2015; OMS, 2010).

Acero and Vasquez (2007) reported in an analysis done of the consultation cases attended in the Child Psychiatry Department of Hospital de la Misericordia in Bogota, that about 20% of children worldwide suffer from a disabling mental illness and that anxiety disorders are included among the most important.

Despite the data obtained from local and global sources and from the report by Acero and Vasquez, the literature on obsessive compulsive disorder in Colombia does not report studies on adults or children with this condition. Latin America lacks studies to address the phenomenon based on clinical aspects, endophenotypes and related genetic characteristics associated with the disorder, which have been reported in the scientific the literature of other areas. Therefore, this study is aimed at determining if the five analyzed polymorphisms and the four neuropsychological tests for executive functions were associated with the presence of the obsessive compulsive disorder in this sample of Colombian children, since the information obtained here is internationally relevant as the reported clinical and biological tests that have generated conflicting results regarding the study of OCD are put to the test.

## Material and Methods

### Participants

For genetic analysis, a total of 63 patients with an average age of 13 were analyzed; to study the relation between polymorphisms and neuropsychology, a subsample of 33 patients and 31 controls was used. The cases were obtained from Hospital de la Misericordia in Bogota, based on the following inclusion criteria: being under 18 years old at the moment of the diagnosis, Colombian nationality, OCD diagnosed and being willing to participate in the study. The inclusion criteria for the 65 control group included: being within the same age range of the patients, no behavioral disorders, no psychiatric medication being administered and a score below 25 points in the Scared test. This study excluded conditions like mental retardation, neurological or metabolic disorders and substance dependence.

The investigation was carried out in accordance with the Declaration of Helsinki 2013, the study design was reviewed by the ethics committee of Universidad Nacional de Colombia-School of Medicine (approval certificate #12) and informed consent and assent of the participants was obtained after the nature of the procedures had been fully explained.

**Table 1.** General characteristics of patients and controls

| Group | Gender |  | Average age | N |
| --- | --- | --- | --- | --- |
|  | Female % | Male % |  |  |
| Patients | 58.2 | 42.8 | 13 | 63 |
| Controls | 53.63 | 46.37 | 10.7 | 65 |

### Procedure

#### Clinical analysis

Each participant and their respective legal guardian were interviewed based on the DSM-IV-TR parameters. Y-BOCS scale and SCARED test were applied to both patients and controls.

#### Instruments

The Stroop test with colors and words, second edition (Golden, 1999) was applied to measure response inhibition, as well as the Trail Making Test (TMT) (Tombaugh, 2004) to test motor skills and spatial monitoring, the Wisconsin Card Sorting Test (WCST) (Heaton, Chelune, Talley, Kay and Curtiss, 2001) to assess behavioral flexibility, and the Tower of London (Injonque and Burin, 2008) as a task to evaluate planning and decision-making.

#### Genetic analysis

DNA was isolated from blood or saliva samples using the Zymo Research^®^ extraction kit and SNPs were genotyped using three different forms of identification. Conventional PCR was performed for 5-HTTLPR, 5HT2A. The bidirectional PCR methodology proposed by Q. Liu, Thorland, Heit, & Sommer, (1997) was followed for the Enzyme COMT1 polymorphism (polymorphism Val158Met). In order to increase specificity, the mismatch technique described by Ayyadevara, Thaden, & Shmookler Reis (2000) and Stadhouders et al., (2010) was used for MAO-A polymorphisms and glutamate transporter. PCR conditions can be seen in Table 2.

**Table 2.** PCR conditions for the thermocycler

|  | SNP | HTTLPR | HT2A | COMT | SLC1A1 | MAO-A |
| --- | --- | --- | --- | --- | --- | --- |
| **Stage** |  |  |  |  |  |  |
| **Denaturation** | x 1 | 94° x 3' | 94° x 3' | 94° x 3' | 95° x 4' | 95° x 4' |
| **Denaturation** |  | 94° x 15" | 94° x 15" | 94° x 15" | 95° x 45" | 95° x 45" |
| **Annealing** | x 35 | 68° x 30" | 65° x 30" | 65° x 30" | 57° x 45" | 58°/52° x 30"* |
| **Extension** |  | 72° x 2' | 72° x 2' | 72° x 2' | 72° x 1' | 72° x 1' |
| **Extension** | x 1 | 72° x 5' | 72° x 5' | 72° x 3' | 72° x 10' | 72° x 10' |
| **Storage** | x 1 | 8 x ∞ | 8 x ∞ | 8 x ∞ | 8 x ∞ | 8 x ∞ |

\* Temperature for first FW1 and 2 respectively

#### Statistical analysis

Statistical analyzes were performed with the PLINK (Purcel et al., 2007) software, using *χ*2 to observe differences between patients and controls; family-based association testing, a linkage disequilibrium (LD) test was performed, as well as a Hardy-Weinberg Equilibrium (HWE) analysis. The Odds ratios (ORs) were calculated with a 95% level of confidence intervals and a Bonferroni correction at 5%. Simple and generalized linear regression models were applied to prove whether there were differences between the results of the tests based on the genotypes, using diagnosis as an auxiliary variable (Core Team, r2015; Dobson, & Barnett, 2008; McCullagh, & Nelder, 1989.) A regression analysis between genotypes and sub-scores on neuropsychological tests was performed for the 33 patients subsample and the 31 controls. For the WCST, a Poisson regression analysis for three of the sub-scores of the test (correct answers, perseveration and conceptualization) was performed; for Stroop, interference scores for TMT (time and number of errors) and for Tower of London (number of trials and runtime) were considered.

## Results

No significant deviations of HWE (p < .05) were found for any of the five polymorphisms. Regarding the association of genetic markers with the presence of the disorder, genes with higher risk levels related to the disorder were associated with SLC1A1 (OR = 1.46) and COMT (OR = 1.487) genes; however, this risk was not significant (p = .1905 and .1358). A TDT analysis was conducted for families but it was not possible to demonstrate a preferential allelic transmission for any of the studied polymorphisms (see Tables 3 and 4.)

**Table 3.** Association analysis for cases and controls

| Gene | A1 | F_A | F_U | A2 | CHISQ | P | OR |
| --- | --- | --- | --- | --- | --- | --- | --- |
| SLC1A1 | G | 0.3083 | 0.2339 | A | 1.714 | 0.1905 | 1.46 |
| 5HT2A1438 | A | 0.4417 | 0.4907 | G | 0.5503 | 0.4582 | 0.8209 |
| HTTLPR | A | 0.5 | 0.5159 | G | 0.063 | 0.8018 | 0.9385 |
| COMTVal158Met | A | 0.4113 | 0.3197 | G | 2.225 | 0.1358 | 1.487 |
| MAOA | G | 0.3118 | 0.2824 | T | 0.1845 | 0.6675 | 1.152 |

A1: Minor allele name (based on whole sample), F\_A: Frequency of this allele in cases, F\_U: Frequency of this allele in controls, A2: Major allele name, CHISQ: chi-square test, P: p-value, OR: Estimated odds ratio

**Table 4.** Family association analysis (TDT)

| Gene | A1 | A2 | T | U | OR | CHISQ | P |
| --- | --- | --- | --- | --- | --- | --- | --- |
| SLC1A1 | G | A | 13 | 14 | 0.9286 | 0.03704 | 0.8474 |
| 5HT2A1438 | A | G | 16 | 16 | 1 | 0 | 1 |
| HTTLPR | A | G | 12 | 18 | 0.6667 | 1.2 | 0.2733 |
| COMTVal158Met | A | G | 23 | 17 | 1.353 | 0.9 | 0.3428 |
| MAOA | G | T | 6 | 12 | 0.5 | 2 | 0.1573 |

A1: Minor allele code, A2: Major allele code, T: Transmitted minor allele count, U:
Untransmitted allele count

WCST results showed differences between cases and controls for the conceptualization score and a tendency towards correct answers and perseverance. During the Stroop test, differences for the interference variable, in which a significant increase of .09 points for patients (p = .041) was reported, were found. In the other TMT and Tower of London sub-scores, no differences between the groups were found.

**Table 5.** Expected differences in WCST.

| SUB-SCORE WCST | EXPECTED SCORE | P VALUE |
| --- | --- | --- |
| Correct answers | +0.084 | 0.062 |
| Perseveration | -4.7 | 0.057 |
| Conceptualization | +14 | 0.023* |

Table 6 shows the regression analysis in which the difference between a reference genotype and a comparison genotype were compared against the sub-scores of the tests. For regression analysis using diagnosis as an auxiliary variable, associations between test sub-scores and HT2A, SLC1A1 and MAO-A genotypes were found. For perseveration score, it was found that the GG genotype of SLC1A1 is associated with greater perseveration (p = .002) and lower conceptualization. Stroop test only presented significant differences between the homozygous for COMT gene (p = .0003), while no significant differences between genotypes were found for the number of trials and perseveration number.

**Table 6.**
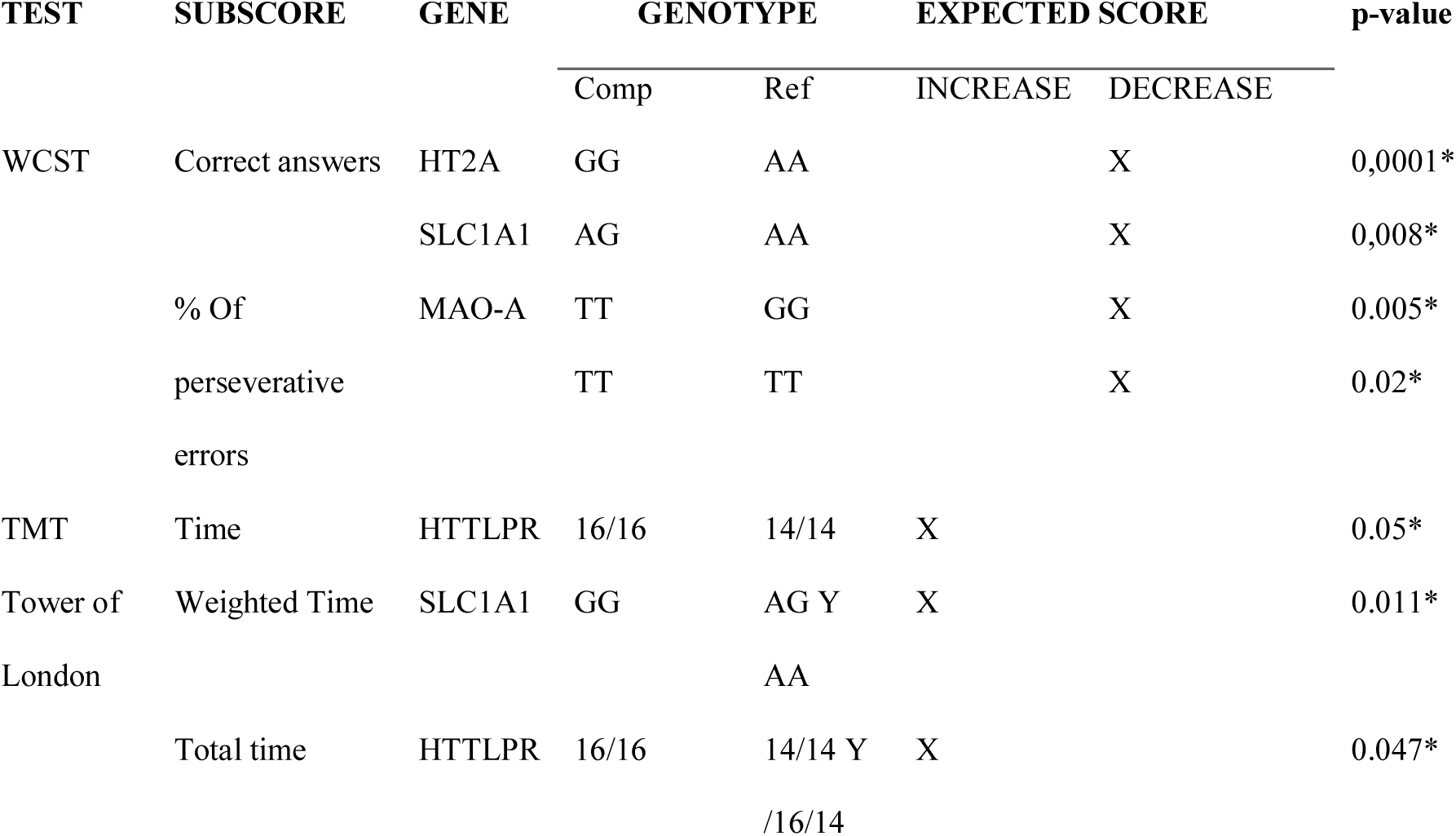
Expected genotype-related differences.

## Discussion

The studies that aim at describing the genetic basis of psychiatric disorders have limited success (Chamberlain and Menzies, 2009) due to several reasons; on the one hand, there is an inherent imprecision in the diagnosis of psychiatric illnesses since patients who may be suffering from a heterogeneous group of disorders are included in the same group. On the other hand, the complexity of the brain as an organ must be considered.

This sample did not find that HTTLPR, HT2A, SLC1A1, COMT and MAO-A polymorphisms were associated with the risk of suffering from the disorder or with protection against it. In this regard, several aspects should be considered, among them, and one of the most important, the fact that genetic association studies reported in the literature show contradictory results for OCD in children and adolescents. It is also worth mentioning that the findings obtained from the GWAS of StewartYu, et al. (2013) and Mattheisen et al. (2015) showed that those polymorphisms more likely to be associated with OCD are not reported in this study. For further genetic studies, considering the genes reported by GWAS to observe the behavior of these genes in a Colombian population is strongly recommended.

A better characterization of the disorder was sought through the use of neuropsychological tests for executive functions in this work and in the work of (Chamberlain & Menzies, 2009; Gottesman & Gould, 2003; Menzies et al., 2007; Viswanath et al., 2009.) TMT, Stroop, Tower of London and WCST were applied, and the latter proved to be the most sensitive one to compare groups. In neuropsychology, results are contradictory for items like flexibility, verbal fluency and decision making tasks (Abramovitch et al., 2013; Flessner et al., 2010; Kuelz et al., 2004; Olley et al., 2007.) For Lewin et al. (2014), neuropsychological studies in children with OCD fail despite efforts on small samples and because test batteries are limited, making the analysis of results difficult. In order to improve the discrimination of cognitive analysis, more complete batteries, in which an IQ test and other motor tests are included to investigate the role of basal ganglia, subscales of intelligence, memory and language, are recommended. These tests should also be made to close relatives of cases and controls to determine whether they function as endophenotypes of the disorder and involve them as a part of the input given to health professionals, professors and caregivers for detecting, treating and follow-up of the functioning of daily tasks performed by children with OCD. The use of innovative testing and the creation of specific neuropsychological tests for the disorder have been proposed (Olley et al., 2007; Lewin et al, 2014.)

The analysis of the relationship between alleles and test scores shows a possible association between cognitive performance and certain genotype setting. This finding should be replicated in a broader sample to deepen the analysis of genetic components that influence the performance of tasks associated with executive functions, independently of the disorder.

Finally, a more complete analysis of sociodemographic characteristics and clinical history of patients is advised. Psychiatric disorders are the result of a different brain development compared to what is considered to be normal (Kofink et al., 2013; Schmitt, Malchow et al., 2014.) Changes and factors associated with the early-onset of psychiatric disorders are complex and, in many cases, little obvious, so further research on the characteristics associated with disorders beyond the symptoms reported in the diagnostic manual must be performed.

